# Preservation of neural synchrony at peak alpha frequency via global synaptic scaling compensates for white matter structural decline over adult lifespan

**DOI:** 10.1101/2021.10.24.465613

**Authors:** Anagh Pathak, Vivek Sharma, Dipanjan Roy, Arpan Banerjee

## Abstract

We propose that preservation of functional integration, estimated from measures of neural synchrony, is a key neurocompensatory mechanism associated with healthy human ageing. To support this proposal, we demonstrate how phase-locking at peak alpha frequency from Magnetoencephalography (MEG) data is invariant over lifespan in a large cohort of human participants, aged 18-88 years. Using empirically derived connection topologies from diffusion tensor imaging (DTI) data, we create an in-silico model of whole-brain alpha dynamics. We show that enhancing inter-areal coupling can cancel the effect of increased axonal transmission delay associated with age-related degeneration of white matter tracts and thus, preserve neural synchrony. Together with analytical solutions for non-biological all-to-all connection scenarios, our model establishes the theoretical principles by which frequency slowing with age, frequently observed in the alpha band in diverse populations, can be viewed as an epiphenomenon of the underlying neurocompensatory mechanisms.

## Introduction

Despite remarkable progress in human neurophysiological and neuroimaging research, a comprehensive theory of brain aging that connects functional neuromarkers with structural constraints and their behavioral ramifications remains elusive. A persistent debate in the field of aging neuroscience centers around the question of whether functional neuromarkers of aging indicate a gradual decay of brain architecture or the presence of compensatory reorganization mechanisms that counteract the deleterious effects of structural loss Park and Reuter-Lorenz [2009], Reuter-Lorenz and Park [2014]. According to the latter view, the brain, being a dynamic and adaptive system can preserve biologically crucial parameters in the face of continual structural decline with age Park and Reuter-Lorenz [2009]. Working point of the system can be homeostatically regulated through plasticity mechanisms that endow the brain with a vast set of dynamical configurations. Loss of one structural component can be compensated for, by an appropriate reconfiguration of system parameters Marder and Goaillard [2006], Turrigiano [1999], Gray and Barnes [2015]. However, the task of classifying specific age-related changes as either adverse or compensatory is complicated by the enormous complexity of brain dynamics Gray and Barnes [2015], Naik et al. [2017].

A case in point is the slowing down of peak alpha frequency (PAF) Babiloni et al. [2006a], Sahoo et al. [2020], a well-documented change in ongoing electro/magneto-encephalographic (EEG/MEG) signals with age. Studies have identified white-matter as a potential locus for resting-state alpha disruption Hindriks et al. [2015], Bells et al. [2017], Minami et al. [2020], Nunez and Srinivasan [2014]. Brain-wide alpha activity is coordinated by and propagates along white-matter fibers Hindriks et al. [2015], Nunez and Srinivasan [2014] that connect spatially distant brain regions. White matter fibres consist of myelinated axons which undergo multiple cycles of repair throughout normal ageing. However, axonal conduction speeds are only partially restored by remyelination, as remyelinated axons possess shorter internodes as compared to developmentally myelinated axons Peters [2009], Scurfield and Latimer [2018]. Reduced conduction speeds along white-matter tracts predict slower network frequencies and impaired synchronization in network models of large-scale brain dynamics Niebur et al. [1991], Pajevic et al. [2014], Petkoski et al. [2018]. Left unchecked, progressive reduction in conduction velocity with age may lead to a complete breakdown of synchrony in crucial brain circuits that subserve normal cognitive processes Nunez and Srinivasan [2014], Sadaghiani et al. [2012]. Hence, from a systems-level view there must exist compensatory mechanisms that arrest functional degeneration.

This article hypothesizes that neural architecture can prevent functional degradation with age by reconfiguring itself in response to the age-related increase in axonal transmission delays. To validate this hypothesis, we estimated a set of measures of neural synchrony— Phase Locking Value (PLV) and Phase Lagged Index (PLI)— in resting-state MEG data obtained from participants aged 18-88 at the Cambridge Centre for Ageing and Neuroscience(Cam-CAN) Shafto et al. [2014]. These measures have been identified as key metrics that define functional integration in brain networks by several researchers Lachaux et al. [1999], Aydore et al. [2013], Cohen [2014]. We find that phase locking (using both PLV and PLI) remains preserved at the IPAF which slows down with age, thus favoring a compensatory role of functional integration during ageing process. Seeking mechanistic insights into this process, we investigate the relationship between IPAF and phase-locking by constructing an in-silico whole-brain model (WBM) of neural coordination. The WBM consisted of coupled differential equations, modeling the phase of autonomous alpha oscillators (Kuramoto model Kuramoto [2003]) at nodes chosen from standardized anatomical parcellations of the human brain Cabral et al. [2011]. White matter properties obtained from diffusion tensor imaging (DTI) data, namely- inter-areal connection strength and transmission delays were varied to study the relationship of steady-state phase-locking and network frequency in the alpha band. The complex network obtained from DTI derived topology and a simplified all-to-all network tractable by mathematical analysis showcased the emergence of a complex interplay between transmission delays and neural coupling as key detyerminants of phase locking and network frequency. Numerically, we explored the regimes of maximal metstability Deco and Kringelbach [2016], Naik et al. [2017], and show how such parameter regimes support the preservation of functional connectivity across lifespan.

## Results

### Alpha phase locking is preserved at the IPAF across age

MEG recordings from the Cam-CAN lifespan cohort (age range 18-88 years) was used to understand the relationship of network frequency and functional connectivity via phase locking measures across age. Source localization was performed on resting-state MEG data from randomly chosen 200 participants in the entire age range from the original sample size of 650 human participants using sLORETA Pascual-Marqui [2002] as implemented by MNE-Python toolbox Gramfort et al. [2014]. After computing the source time series at predefined anatomical parcellations (Desikan et al. [2006]), 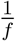 fluctuations were removed using an automated algorithm Donoghue et al. [2020] for identification of individual peak alpha frequency in each participant (IPAF). This was deemed necessary because 1/f features in EEG have been shown to vary with age Voytek et al. [2015]. Out of 200, 160 participants possessing a distinct alpha peak in at least 30 ROIs(out of 68) were included in the final analysis. For these participants, mean peak alpha frequency (average peak alpha across ROIs) was found to significantly reduce with age (*r* = −0.4, *p* < 0.0001) (**Figure 2**), similar to patterns reported in earlier studies Sahoo et al. [2020].

Next, for each participant we computed the Phase locking value (PLV) at the IPAF with a bandwidth of 4*Hz* between all ROI pairs. PLVs were averaged across all possible pairs to obtain one PLV for each participant. For comparison, we also estimated PLV at two other frequency bands - lower alpha (LA,6-10Hz) and upper alpha (UA, 10-14Hz). An additional measure of phase locking — Phase Locking Index (PLI), was estimated to rule out the effect of volume conduction manifesting as zero-lag phase synchronization (**Figure 1**). Correlation analysis between age and band-specific phase locking revealed that both PLV and PLI increased with age in LA band (*r_PLV_* = 0.21, *p_PLV_* < 0.001, *r_PLI_* = 0.29, *p_PLI_* < 0.001); PLV and PLI decreased with age in the UA band(*r_PLV_* = *−*0.38, *p_PLV_* < 0.0001, *r_PLI_* = *−*0.3, *p_PLI_* < 0.001). In contrast, PLV and PLI remained invariant with age at the IPAF centered band (*r_PLV_* = − 0.1, *p_PLV_* = 0.16, *r_PLI_* = 0.09, *p_PLI_* = 0.29) (**Figure 2**). Surrogate distributions, obtained by randomly shuffling resting state epochs indicated that PLI values in the IPAF centered band were significantly higher than what would be expected by chance (*p* < 0.01) (**Figure 2**). Taken together, the results indicate that phase locking is preserved at the IPAF, which slows with age(**Figure 2**). Findings were also replicated at the sensor level (N= 650, see Supplementary material).

**Figure 1.**
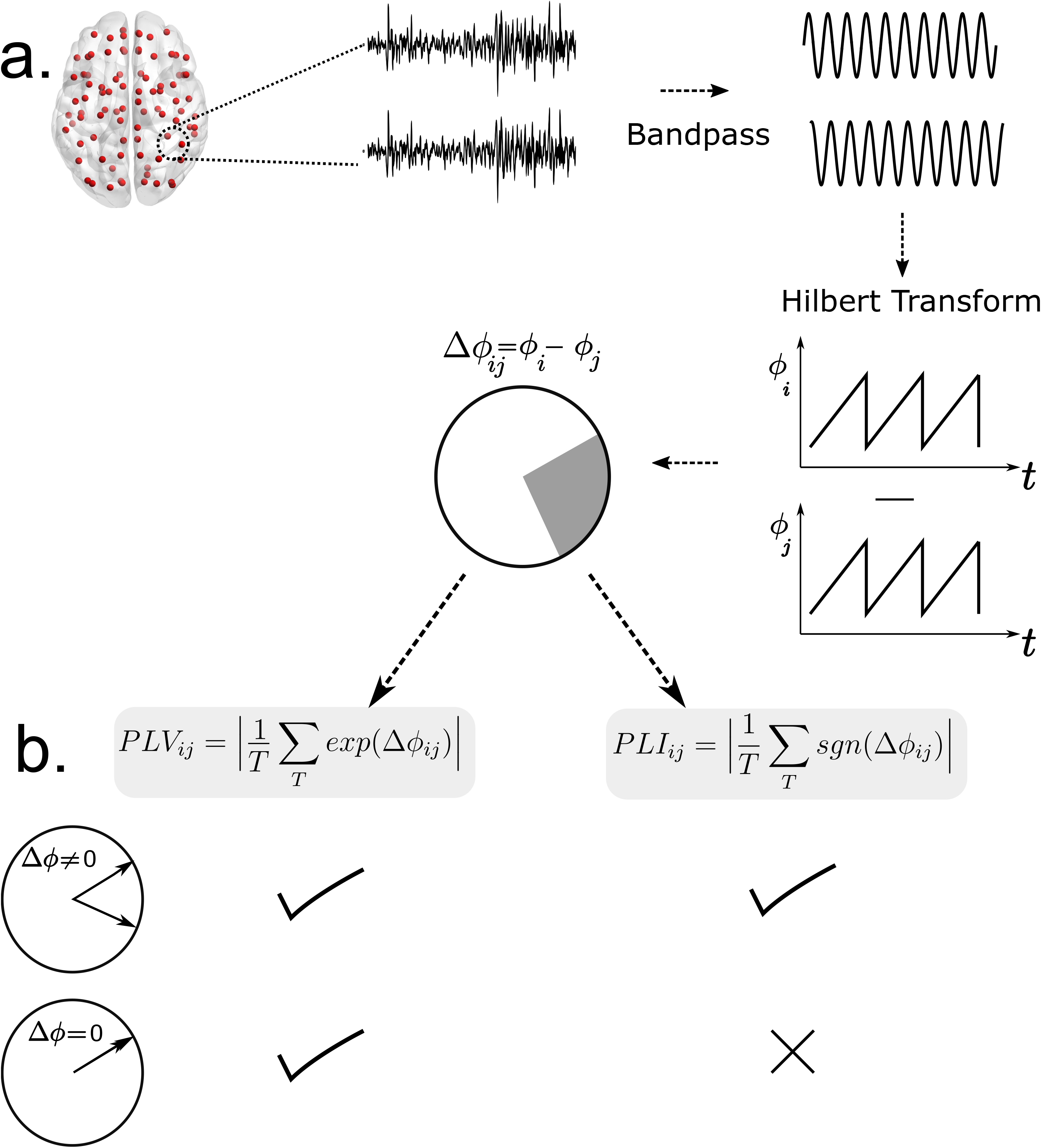
Pipeline for estimation of phase locking via PLV and PLI. **a)** Source localized MEG signals are bandpass filtered to extract signals in specific frequency bands. Hilbert transform is used to extract instantaneous phase time series. Phases at each timepoint are projected onto a unit circle. **b)** Mathematical expressions for the estimation of PLV and PLI. PLV measures zero-phase lags whereas PLI discounts zero phase lags. This property makes PLI resilient to volume conduction/field spread artifacts that manifest as zero-phase lag correlations.

**Figure 2.**
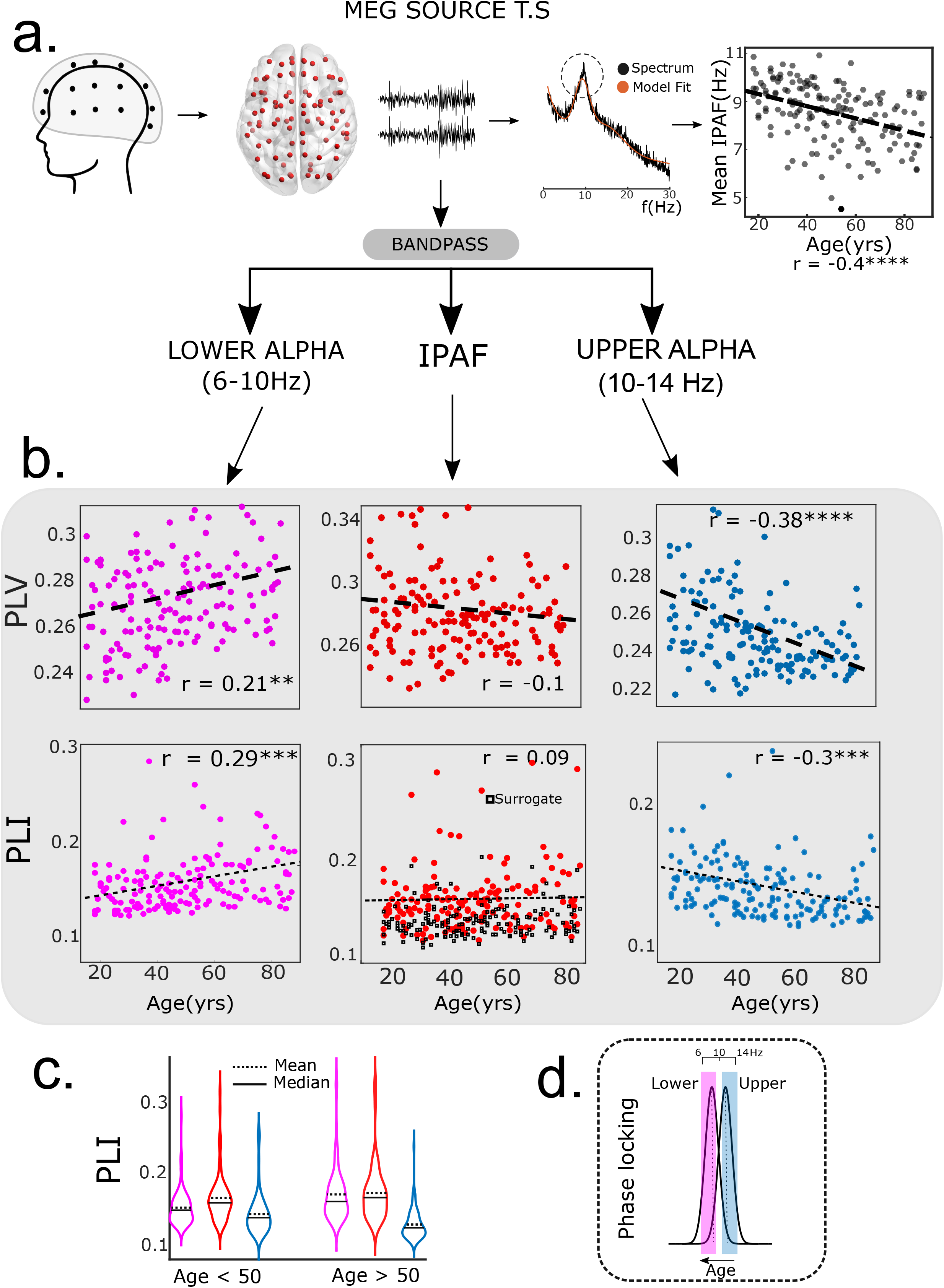
Phase locking in the alpha band. **a)** Overview of analysis pipeline: rsMEG sensor space data was source localized(sLORETA) and projected to a standard parcellation(Desikan-Killainy). PSD for each ROI was extracted using welch method and modeled as a linear superposition of periodic and aperiodic components. Peak frequency was extracted for each brain region and averaged across ROIs to obtain a single mean peak alpha frequency for each subject. Mean peak alpha frequency was found to be negatively correlated with age. Subsequently, phase locking was estimated for each subject using both PLV and PLI. **b)** Phase locking value (PLV) and Phase locking index(PLI) estimated for three frequency bands-LA(6-10Hz), IPAF(IPAF-2–IPAF+2) and UA(10-14Hz). **c)** PLI values for LA, IPAF and UA band. **d)** Schematic: PLV, PLI analysis suggests frequency re-organization that preserves alpha phase locking at reduced peak frequencies.

### Conduction delays and coupling modulate network frequency and synchrony in an idealized neural network with all-to-all connections

We motivate a theoretical understanding of how oscillatory frequency and network synchronization are modulated via connection properties by considering a network of N, Kuramoto phase-oscillators Kuramoto [2003]. Oscillators interact with one another according to the following equation-

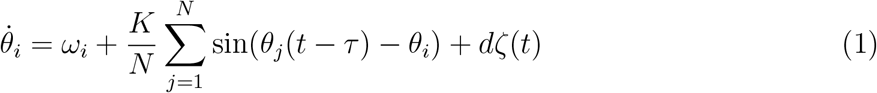

where, *θ* and *ω* are the phase and natural frequency of each oscillator. *K* and *τ* specify average coupling strength and transmission delay between any two nodes respectively. Natural frequencies are derived from a symmetrical distribution centered at *μ*. In the most general case, the system is supplied with zero mean Gaussian noise process *ζ*(*t*), with a standard deviation *d*. The network is composed of N oscillators connected according to an all-to-all topology. The order parameter (r) indexes the degree of phase-synchronization in the network

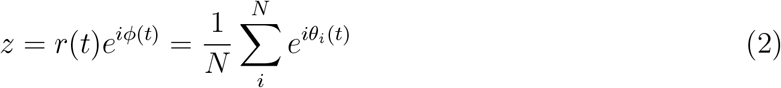

such that 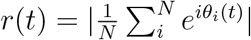, with *r* = 0 corresponding to incoherence and *r* = 1 to complete synchronization and *z* is a complex valued function tracking the global phase synchronization in the network. For smaller coupling values, the incoherent state is stable. The incoherent state loses stability at a critical value of coupling(*K_c_*), giving rise to a partially synchronized regime. In the absence of conduction delays (*τ* = 0), the network of oscillators synchronize at the center frequency (*μ* = 10*Hz*) for *K > K_c_*. However, for *τ* ≠ 0, the synchronization frequency(Ω) is different from the center frequency of the distribution of natural frequencies (*μ*)(**Figure 3**) Niebur et al. [1991]. Specifically, we observe Ω < *μ* for the all-to-all coupled network considered here.

**Figure 3.**
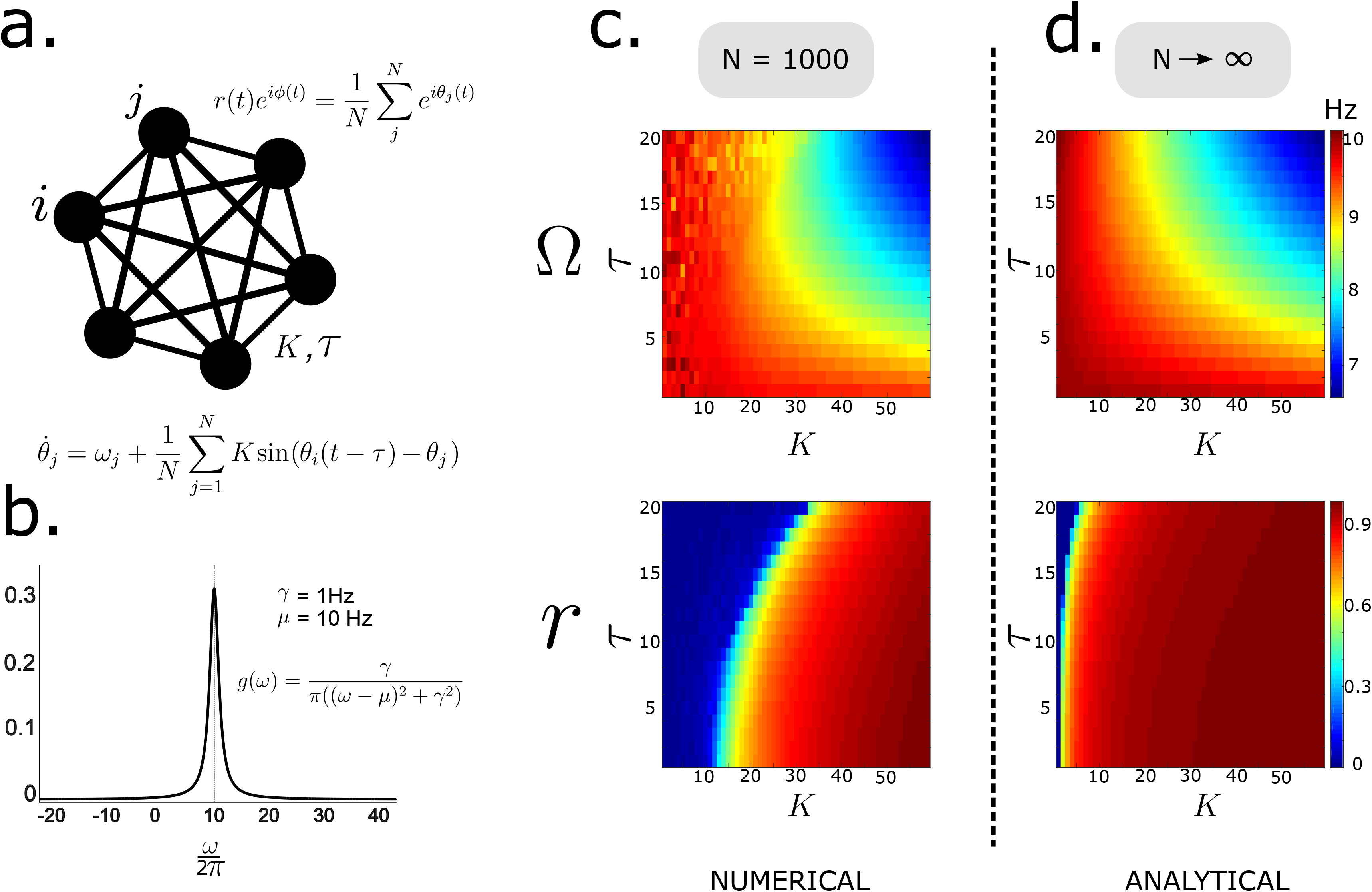
Phase dynamics in an idealized network. **a)** Fully recurrent network of phase oscillators is considered. **b)** Natural frequencies of oscillators are drawn from a Lorentzian distribution with, *γ* = 1*Hz*, *μ* = 10*Hz*. **c)** Steady state synchronization frequency and order parameter for *N* = 1000 oscillators (*d* = 0), obtained by numerical simulations. Delays were varied between 0 – 20*ms*. **d)** Analytical expressions for synchronization frequency and order parameter derived by reducing the high dimensional system through the Ott-Antonsen method. Network frequency and order parameter are modulated by coupling and delay.

For non-zero delays, the network exhibits multistable states, such that the system can reside in multiple synchronized regimes (see analytical solution below), each associated with a different synchronization frequency Niebur et al. [1991], Yeung and Strogatz [1999]. Heatmap in **Figure 3c**, shows the relationship of steady state collective frequency of the network (colour) for different (*K, τ*). For smaller values of conduction delays, the incoherent network has an average frequency close to the mean of the distribution of natural frequencies (*μ* = 10*Hz*). However, for longer delays, the collective synchronization frequency shows significant suppression.

The order parameter can be shown to evolve via a low-dimensional system of global synchronization manifold under a set of simplifying assumptions Ott and Antonsen [2008]. Expressions for steady state synchronization frequency and order parameter were obtained from the low dimensional system (see Analytical solution) and compared with parameter space obtained from numerical simulations 1.

#### Analytical Solution relating synchronization frequency and conduction delays for a reduced system

We derive analytical expressions for the synchronization frequency (Ω) and steady state order parameter (r) for the case of a fully recurrent network of phase oscillators (*N* → ∞), connected to each other via coupling *K* subject to delay *τ*. For simplicity, we consider the noiseless case (*d* = 0). The natural frequencies of oscillators *ω_j_* are derived from a *Lorentzian* distribution given by

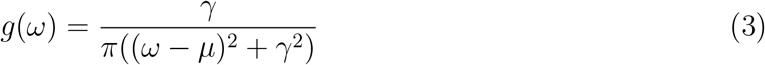

Ott and AntonsenOtt and Antonsen [2008] showed that the macroscopic dynamics corresponding to equation 1 follows a low dimensional ODE given by-

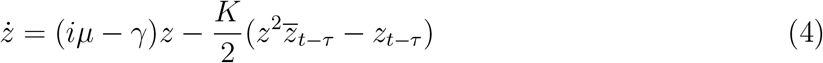

For details of this step please refer to the Supplementary Material or Ott and Antonsen [2008]. We demand steady-state solutions of the form Ott and Antonsen [2008]

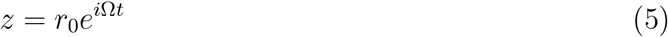

where *r*_0_ and Ω are the steady state order parameter and synchronization frequency respectively. Equations 4 and 5 lead to

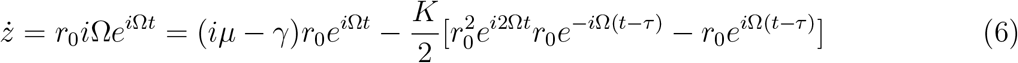

*r*_0_ = 0 (incoherent solution) is a trivial solution of equation 6 for all *K, τ, γ*. In order to explore coherent solutions we equate the real and imaginary parts on both sides leading to the following transcendental equations

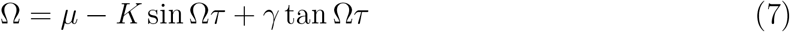

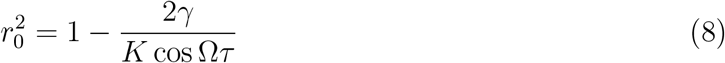

The requirement 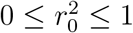 yields the following condition for the existence of coherent solutions

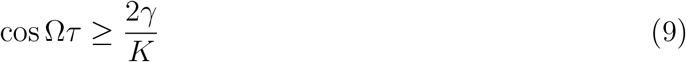

Equation 7 and 8 suggest a mechanism through which frequency and order parameter are modulated as a function of coupling and delay for given *γ* and *μ*. The transcendental equation 7 can be approximately solved by performing Taylor series expansions, 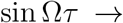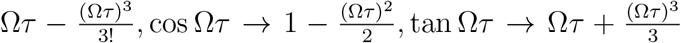. Considering the first two terms of the Taylor series the transcendental equation 7 can be simplified to

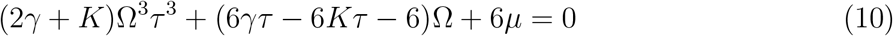

and the constraint in 9 can be further approximated to

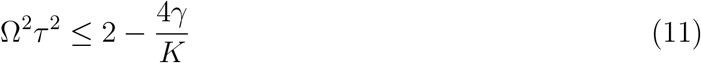

Subsequently, equation 10 is solved numerically by using the MATLAB routine *fsolve* for the parameter space constrained by the condition 11. The solution shows excellent agreement with numerical solutions with N = 1000, as shown in **Figure 3**. Most importantly for our study, we find that reductions in network synchrony due to increased conduction delays can be offset by concomitant changes in coupling.

### Reduced network synchrony due to lowered conduction speeds can be rescued by global scaling of connection strength

In-silico modelling of brain dynamics was used to study whether alpha dynamics on white-matter network topology retains the key features exhibited by idealized networks for which analytical relationships between network frequency and neuronal coupling are derived in the previous section. Kuramoto phase-oscillators Kuramoto [2003] are placed at anatomical landmarks using the Desikan-Killiany atlas Desikan et al. [2006]. For specifying white-matter connectivity, we used DTI adjacency matrices from a separate dataset of healthy subjects, as described in Abeysuriya et al. [2018]. Conduction delays between all node pairs were estimated by scaling inter-node Euclidean distances by a fixed cortical conduction velocity *v*, 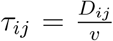. The biological scenario differs from the idealized network in three key aspects-1. Topology, 2. Distribution of natural frequencies and 3. Introduction of distance dependent delays. Oscillators are set up to interact with one another according to the following equation Kuramoto [2003]

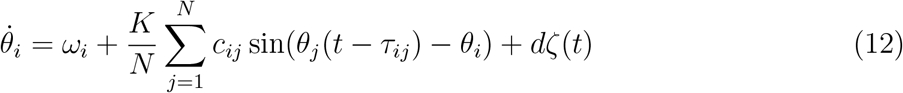

where, the term *c_ij_* introduces heterogeneous connection weights derived from normalized measures of fibre densities. Similar to the idealized case, network frequency and phase locking values were obtained by varying cortical conduction velocity(v) and global scaling parameter(K). Cortical conduction velocity was varied in the range of 1 30*m/s*, in line with previous experimental reports Swadlow [1982]. Altering the conduction velocity changes the distribution of distance dependent transmission delays, whereas changing *K* is analogous to synaptic scaling. Following Gollo et al. [2017], Roberts et al. [2019], natural frequencies (*ω_i_*’s) are distributed across ROIs based on node strength (*ω_max_* = 12*Hz, ω_min_* = 8*Hz*, *μ* = 11.06*Hz*) (**Figure 4,b**, see Methods for selection of natural frequencies).

**Figure 4.**
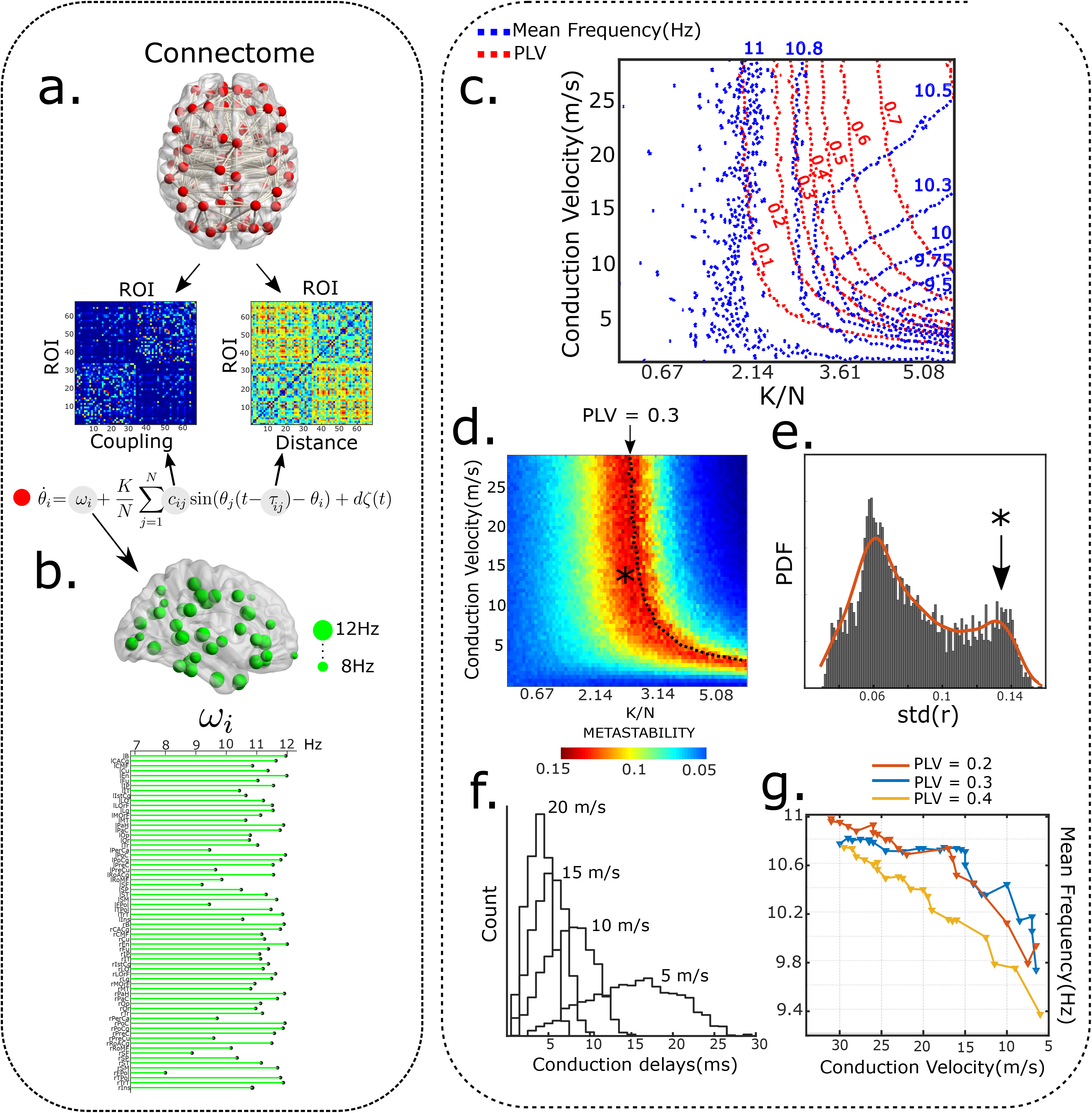
Large scale alpha phase locking. **a)** Model overview-(Above) DTI connectivity and distribution of inter-node distances. (Below)Equations governing node dynamics. **b)** Distribution of natural frequencies. (Above)Green spheres represent magnitude of natural frequency. (Below) ROI wise distribution of natural frequencies. **c)** Contour plot indicating isolines for mean frequency(blue) and PLV(red) as a function of global coupling and conduction velocity, Noise amplitude(d) = 3, *ω_max_* = 12*Hz*, *ω_min_* = 8*Hz*. PLV and PAF remain constant along isolines. **d)** Metastability measured as the standard deviation of the order parameter plotted as a function of conduction velocity and global coupling. Dotted line indicates PLV isoline. **e)** Distribution of metastability distribution. Red region in heatmap corresponds to second mode of the gaussian. **f)** Distribution of conduction delays(in ms), for conduction velocity=5,10,15.20 m/s **g)** Frequency depression along isolines corresponding to PLV = 0.2, 0.3 and 0.4.

Similar to observations in idealized network topology with all-to-all connections, network synchrony is modulated by the combined influence of conduction velocity and global gain parameter (K). Broadly, the system exhibits lower levels of synchrony for very weak coupling and low conduction velocity (**Figure 4**). On the other hand, larger coupling and conduction velocity lead to hyper-synchronous states. Somewhere in between lies the partially synchronized state, characterized by high temporal variability in the Kuramoto order parameter. Such maximally metastable regimes are thought to underlie resting state dynamics Deco and Kringelbach [2016], Naskar et al. [2021]. Accordingly, we restrict our attention to the metastable regime while considering age-related reduction in conduction velocity. Distribution of metastability values clearly delineates the metastable regime as a distinct mode (**Figure 4,e**).

To better visualize the relationship of network frequency and phase locking, we plot contour lines. Compensatory balancing of phase locking corresponds to a traversal along the PLV contour lines (**Figure 4,c**). We observe a robust reduction in mean network frequency along PLV contour lines in the metastable regime (**Figure 4,c,d**). In contrast, a vertical descent along the y-axis in **Figure 4,a**, that can be interpreted as a passive decline in conduction velocity without compensation, is not accompanied by any significant reduction in network frequency. This can be gauged by the vertical orientation of frequency contours in the metastable regime.

Interestingly, moving along the PLV contour lines not only preserves network synchrony, but also metastability (red region, **Figure 4d**). Therefore, in addition to preserving mean synchrony levels, synaptic scaling also maintains the temporal richness of alpha dynamics that subserves the dynamic repetoire of core brain areas Deco et al. [2017]. A non-compensatory decline in network conduction velocity also raises the possibility of a sudden increase in network frequency owing to a complete breakdown of network synchrony due to high network delays. However, traversing along PLV contour lines assures a monotonic reduction of network frequency, while maintaining synchronicity among brain areas. Model simulations with different noise amplitudes and distributions of natural frequency led to qualitatively similar results(**Supplementary**). To demonstrate the generality of our results, we replicated the analysis using a different connectomic dataset and parcellation scheme(automated anatomical labeling,AAL) as described in Cabral et al. [2014](**Supplementary**).

## Discussion

In this study we propose that functional integration achieved via neural synchrony is a dominant neurocompensatory mechanism that is underway during healthy lifespan ageing. The key entry point that led us towards this understanding is a widely reported phenomenon – age-related decrease in individual peak alpha frequency (IPAF) Babiloni et al. [2006a], Voytek et al. [2015], Scally et al. [2018], Sahoo et al. [2020] which we demonstrate to be the outcome of a dynamic functional compensation process that preserves network synchrony in response to inclement enhancement transmission delays in information propagation stemming from demyelination of axonal tracts as function of age. We test our hypothesis on empirical MEG recordings by estimating measures of network synchrony at the IPAF in both source (**Figure 2**) and sensor levels (Supplementary Material). We validate this hypothesis by employing two complementary measures of phase synchrony-phase locking value (PLV) and phase lag index (PLI) on source-localized MEG data made publicly available by Cam-CAN. While PLV estimates consistency of phase differences, PLI additionally controls for volume conduction by discounting zero-lags Stam et al. [2007]. Through surrogate testing we confirm that PLI values indicate significant phase relationships and are not artifacts of sample size bias Stam et al. [2007]. Our experimental results corroborate the findings of preserved IPAF connectivity observations by Scally and colleagues which were published for a smaller sample size on EEG sensor level data Scally et al. [2018]. Here we also demonstrate the preservation of network synchrony at IPAF in both source and sensor level data, thus confirming and expanding the scope of previous findings Scally et al. [2018]. To further elucidate the mechanistic basis of neurocompensatory mechanisms, we employ computational modeling to gain insights into the dynamic origins of age-associated IPAF slowing. Numerous studies have speculated a prominent role for white-matter fibers in modulating alpha synchrony Bells et al. [2017], Hindriks et al. [2015], Minami et al. [2020]. Therefore, in order to study the relationship of network frequency and synchrony, we reduce large-scale white-matter network to its basic dynamical elements– conduction delays and inter-areal coupling that forms the backbone of a whole brain connectome. Each node in the connectome is considered to be a unit amplitude limit-cycle oscillator (an idealized autonomous oscillator), described by its phase (**Figure 3**). Anatomically, each autonomous alpha oscillator can be identified with a self-sustained thalamo-cortical unit, or alternatively, pacemaker populations such as the infragranular and supragranular layer in V2 and V4 Victor et al. [2011], Bollimunta et al. [2008]. Both numerical and analytical approaches on idealized network with all-to-all connections confirm how changes in average conduction delays may be offset by modifications in coupling, and that frequency slowing is a collateral to this compensation. Next, we extend our model to include network topology estimated from empirical human white-matter connectivity and find robust frequency modulation with average conduction speed change and global coupling. We track trajectories in the global coupling-conduction velocity space that preserve phase locking, restricting our attention to the partially synchronized regime. Our results unequivocally demostrates frequency slowing emerges as the system attempts to maintain phase locking in response to a reduction in conduction velocity by modulation of inter-areal connectivity (**Figure 4**).

As early as the 1950s, Norbert Wiener had hypothesized that independent oscillators with natural frequencies close to 10 Hz interact with one another to shape alpha rhythmicity Wiener [1966], Strogatz [1994]. According to Wiener, alpha activity emerges from frequency pulling between individual alpha oscillators which possess slightly different natural frequencies, such that the system of oscillators “constitute a more accurate oscillator *en masse* than they do singly” Wiener [1966]. This idea is regarded as one of the earliest models of collective dynamics of biological oscillators Strogatz [1994]. In the intervening years, models of collective synchronization have become a mainstay of neuroscience, having been employed to explain diverse phenomenon such as travelling brain waves, fluctuating beta oscillations, fMRI functional connectivity, large scale brain synchronization, myelin plasticity etc. Cabral et al. [2011], Zhang et al. [2018], Breakspear et al. [2010], Petkoski et al. [2018], Noori et al. [2020]. In the present article, we adapt Wiener’s idea of frequency pulling to explain the gradual slowing of alpha frequency with age. Specifically, we show that frequency pulling in the presence of conduction delays, biases the system to synchronize at lower frequencies. Intuitively, the mechanism proposed here is analogous to a group dance, where complicated dance moves are initially practiced slowly, since it is easier to maintain lockstep at lower speeds. Similarly, upscaling inter-areal coupling and inadvertently global synaptic scaling allows for the maintenance of network *lockstep* at slower coordination frequencies.

Similar homeostatic mechanisms that regulate circuit output have been identified elsewhere. For example, neurons in the visual cortex of developing rodents undergo synaptic scaling in response to visual inputs Desai et al. [2002]. Synaptic scaling has also been shown to compensate for neuron number variability in the crustacean stomatogastric ganglion Daur et al. [2012]. Recently, Santin and colleagues elegantly demonstrate how respiratory motor neurons in the bullfrog can dynamically regulate breathing by modifying synaptic strengths after long periods of inactivity Santin et al. [2017]. From the perspective of communication through coherence (CTC) hypothesis Fries [2015], alpha phase-locking constitutes an *information channel*, whereby distant oscillators with slightly different peak frequencies communicate with one another through leader-laggard phase relations. Consistent phase locking, a prerequisite for effective communication across brain regions, entails that oscillators adjust their individual frequencies under the combined influence of coupling and transmission delays. Thus, there emerges a clear relationship between oscillation frequency and phase connectivity. Therefore, our central hypothesis is that frequency shifts with aging need to be understood in the context of homeostatic maintenance of large-scale phase locking. Understanding the precise mechanism of frequency slowing – whether adverse or compensatory – has far-reaching consequences for characterizing age-associated neuropathologies like Dementia and Alzheimer’s disease, which share PAF slowing as a prominent featureLópez-Sanz et al. [2016], Sharma and Nadkarni [2020], Babiloni et al. [2006b]. In the framework proposed here, greater frequency slowing in AD may result from the higher demands placed on compensatory processes by accelerated demyelination. Therefore, our model supports a growing view that suggests a greater role for white-matter abnormality in explaining AD progression Sachdev et al. [2013].

Compensatory models of aging have been proposed to account for the finding that many individuals continue to function remarkably well with age despite significant structural loss. For example, according to the scaffolding theory of aging and cognition(STAC) Park and Reuter-Lorenz [2009], the aging brain can preserve cognitive function in the face of age-associated neural changes like volume shrinkage, white-matter degeneration, cortical thinning, and dopamine depletion by recruiting alternate neural pathways, referred as scaffolds. While the STAC model has succeeded in explaining a number of observations in aging neuroscience at the cognitive level, we still lack a clear understanding of the dynamical principles that facilitate compensation or in other words a physical quantity that remains invariant. Thus, our key empirical finding that preservation of phase locking at PAF gives a novel insight to compensatory processes that are involved by the candidate brain networks. Thus, our model departs from the standard conceptualization of alpha slowing as an adverse outcome of aging. Rather, we recast frequency slowing as a tell-tale signature of neural compensation. Earlier models have conceptualized alpha slowing as a passive process, resulting from the gradual decline of system parameters. For example, Lopes da Silva and colleagues Da Silva et al. [1974] model EEG maturation by using a neural mean field model. Their model consists of two populations of neurons: thalamic and cortical, driven by multiple uncorrelated noise sources. By changing the feedback coupling parameters the authors obtained a family of spectral curves that closely resemble developmental trajectories. However, the model lacks axonal delays, which are known to undergo age-related changes and are held to be major drivers for the evolution of cortical networks. Similarly, Van Albada and colleagues Van Albada et al. [2010] employ a more detailed neural mean field model of the thalamo-cortical system Robinson et al. [2001] to investigate age-associated changes in EEG spectral parameters and found white-matter stabilization and regression to be a major determinant of EEG characteristics across life-span. However, the approach, models the gross EEG spectrum for estimating model parameters, making it hard to dissociate specific mechanisms responsible for age-related changes in narrow-band frequencies. More recently, Bhattacharya et. al. Bhattacharya et al. [2011], used a variant of the Lopes Da Silva model to study slowing of peak alpha oscillations in the context of Alzheimer’s disease, implicating thalamic inhibition as the principal driver of alpha slowing, however, as with the original Lopes Da Silva model, this model does not consider axonal conduction delays. Future efforts can build on the model proposed here to tease out how compensatory processes operate in various other contexts, such as rehabilitation from stroke, recovery from traumatic brain injuries to predict recovery timelines and to detect critical times for intervention.

## Methods

### MEG analysis

#### Data description

Data used in the preparation of this work was obtained from the CamCAN repository (available at http://www.mrc-cbu.cam.ac.uk/datasets/camcan/) Shafto et al. [2014]. Cam-CAN dataset was collected in compliance with the Helsinki Declaration, and has been approved by the local ethics committee, Cambridgeshire 2 Research Ethics Committee (reference:10/H0308/50). For all the subjects, MEG data were collected using a 306-sensor (102 magnetometers and 204 orthogonal planar magnetometers) VectorView MEG System by Elekta Neuromag, Helsinki, located at MRC-CBSU. Data were digitized at 1 kHz with a high pass filter of cutoff 0.03 Hz. Head position was monitored continuously using four Head Position Indicator coils. Horizontal and vertical electrooculogram were recorded using two pairs of bipolar electrodes. One pair of bipolar electrodes were used to record electrocardiogram for pulse-related artifact removal during offline analysis. The data presented here consisted only of resting state, where the subjects sat still with their eyes closed for a minimum duration of 8 min and 40s.

#### Data selection and Source Reconstruction

200 subjects from the original data were randomly selected for further source reconstruction analysis. To ensure homogeneous sampling, we sampled 50 participants from each of the following age-groups— Young(18-34 yrs), Middle Elderly(35-49 yrs), Middle late(50-64)yrs and Old(65-88 yrs). Random selections were repeated until a relatively equal split between genders was obtained across each age-group. Pre-processed MEG data of selected participants was referenced to a standard template(Collins27) Holmes et al. [1998]. MRI segmentation was performed using freesurfer. Boundary element method was used to compute surface triangulation for forward computation Fischl [2012]. Standard low resolution brain electromagnetic tomography(sLORETA), implemented in MNE-python was employed for source estimation Gramfort et al. [2014]. Source time-series were projected to 68 brain parcellations according to the Desikan-Killiany atlas Desikan et al. [2006].

#### IPAF estimation

Time series from each participant were epoched in 5s bins and downsampled to 90 Hz. The influence of EOG and ECG on sensor data was removed for every subject through an automated ICA algorithm Sahoo et al. [2020]. All subsequent analysis was performed on the resulting data. ROI-wise power spectral density was estimated using Welch periodogram method. Spectrum was estimated for each epoch after multiplying time series with a Hanning window. Subject spectrum was obtained by averaging across epochs. Power spectrum of electrophysiological recordings consist of both periodic and aperiodic components Voytek et al. [2015], Thuwal et al. [2021]. In order to remove the influence of aperiodic 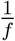 component, the spectrum(P) is modelled as -

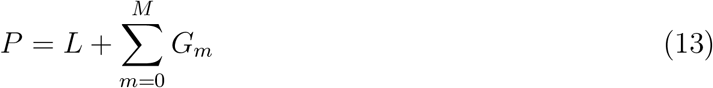

*L*,*G_m_* model the aperiodic and periodic components respectively. *G_m_* is approximated a a *Gaussian* function-

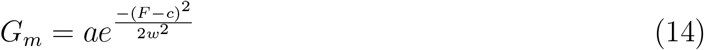

while the aperiodic component(*L*) is modeled in the semi-log power space as-

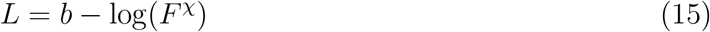

where, *b* is an offset and *χ* is the slope. *a*, *c*, *w*, *b* and *χ* were estimated through an automated model fitting procedure as described in Donoghue et al. [2020]. Model fitting was performed in the 2-20 Hz range, following Tran et al. [2020]. We excluded participants who had distinct peaks in fewer than 30 ROIs in the alpha band(6-14Hz).

#### PLV and PLI

Phase locking was estimated by using PLV and PLI measure. Firstly, each 5s epoch for each participant was bandpass filtered in LA, UA and SSA band. Next, Hilbert transform was performed for each filtered epoch to extract phase time series, *ϕ_a_*(*t*). Phase difference (*ϕ_ab_*(*t*) = *ϕ_a_*(*t*) − *ϕ_b_*(*t*)) was calculated for each ROI pair. PLV and PLI were estimated as Stam et al. [2007]-

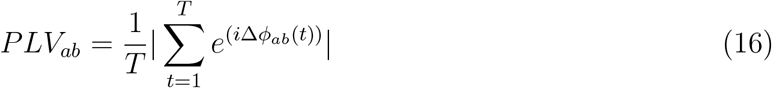

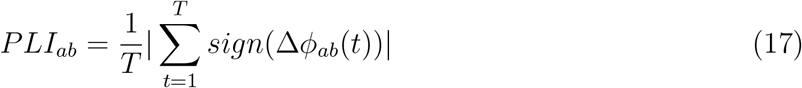

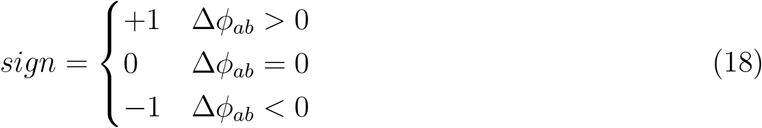

PLV and PLI were averaged across epochs and ROI pairs.

#### Permutation Testing

Pearson’s linear correlation was computed to calculate age-trends for IPAF, PLV and PLI. Surrogate distributions were generated by randomly shuffling variables across 10,000 iterations. Additionally, bootstrapping was performed to ascertain significant PLI in the SSA band. Surrogate distribution for PLI was obtained by randomly shuffling epochs to produce 100 PLI values; p-values were estimated from the resulting distribution.

### Whole-brain model of alpha slowing

In this study we use the Kuramoto model 1 with conduction delays in order to explain brainwide slowing of alpha oscillations with age. We demonstrate frequency slowing on two types of topology-1) Fully recurrent, 2) DTI based structural connectivity.

#### Structural connectivity(DTI)

For our study we use structural connectivity(SC) matrix derived from Human Connectome Project as provided in Abeysuriya et al. [2018]. SC matrices were obtained by performing probabilistic tractography on diffusion MRI data. In short, fibre orientations were calculated from distortion-corrected data, as implemented in FSL. Probtrackx2 was used to detect upto 3 fibre orientations per white-matter voxel. Matrices were reduced to a 68 * 68 scheme, according to the Desikan-Killainy atlas(Desikan et al. [2006]). Adjacency matrices of 40 participants were averaged. Log-transformation was performed to account for algorithmic biases. Conduction delays were obtained by scaling barycentric distances between ROIs by conduction velocity.

#### Natural frequency assignment

Natural frequencies for 68 ROIs were assigned based on anatomical node strengths according to the equation Gollo et al. [2017], Roberts et al. [2019]-

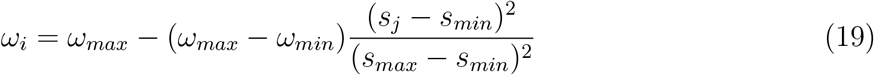

Where, *ω_max_* = 12*Hz* and *ω_min_* = 8*Hz* are the maximum and minimum oscillatory frequencies, specifying the distribution of alpha frequency across ROIs. *s_max_* and *s_min_* are the maximum and minimum strengths respectively.

#### Metastability

Metastability refers to the ability of dynamical systems to flexibly engage and disengage without remaining confined in trivial dynamical configurations such as hyper synchrony or incoherence. According to the communication through coherence(CTC) view, brain areas communicate through state dependent phase coupling Deco and Kringelbach [2016]. In this scheme, resting state dynamics must display maximal variability in phase configurations(i.e maximal metastability). The principle of maximal metastability, combined with realistic values of cortical conduction speeds allow us to demarcate relevant regions for exploration in the conduction speed-coupling space. The standard deviation of the order parameter 2 is regarded as a proxy for metastability Deco and Kringelbach [2016].

#### Numerical Integration

For the recurrent network, system of equations represented by 1 was numerically solved for *N* = 1000 oscillators using the Euler method. Integration time step was kept at *dt* = 0.001*s*. For DTI connectivity conduction delays were assumed to be integer multiples of *dt* to avoid use of computationally intensive interpolation schemes. Noise was supplied to each node by multiplying random normal numbers by the noise amplitude, scaled by 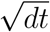. Each simulation was run for 30*s* and first 10*s* were discarded and all subsequent analysis was performed with the resulting signal. Neuroscience gateway platform was used to simulate computationally intensive parameter sweeps Sivagnanam et al. [2013].

## Acknowledgements

We acknowledge the generous support of NBRC Core funds and the Computing facility. For simulations, resources from Neuroscience Gateway Sivagnanam et al. [2013] were used. Data collection and sharing for this project was provided by the Cambridge Centre for Ageing and Neuroscience (Cam-CAN). Cam-CAN was supported by the UK Biotechnology and Biological Sciences Research Council (Grant BB/H008217/1), together with support from the UK Medical Research Council and University of Cambridge, UK. In accordance with the data usage agreement for Cam-CAN dataset, the article has been submitted as open access.

