## Supplementary for "Preservation of neural synchrony at peak alpha frequency via global synaptic scaling compensates for white matter structural decline over adult lifespan"

### SUPPLEMENTARY INFORMATION

#### 1. Frequency suppression for different Noise Amplitudes and Natural Frequencies

In order to establish the generality of our results, we repeated the analysis with different levels of noise inputs and by selecting different natural frequency bands. Just as in the main text, the metastability index, network frequency and phase locking value were estimated by varying conduction velocity(between 1-30m/s) and global scaling parameter(K). The parameter d, which scales noise amplitude, was set to 2,4. In the last row, we show parameter sweeps for the system when natural frequencies are scaled between $\omega_{min} = 6\,Hz$ and $\omega_{max} = 14\,Hz$. The third column demonstrates compensatory reduction of peak frequencies for different values of PLV.

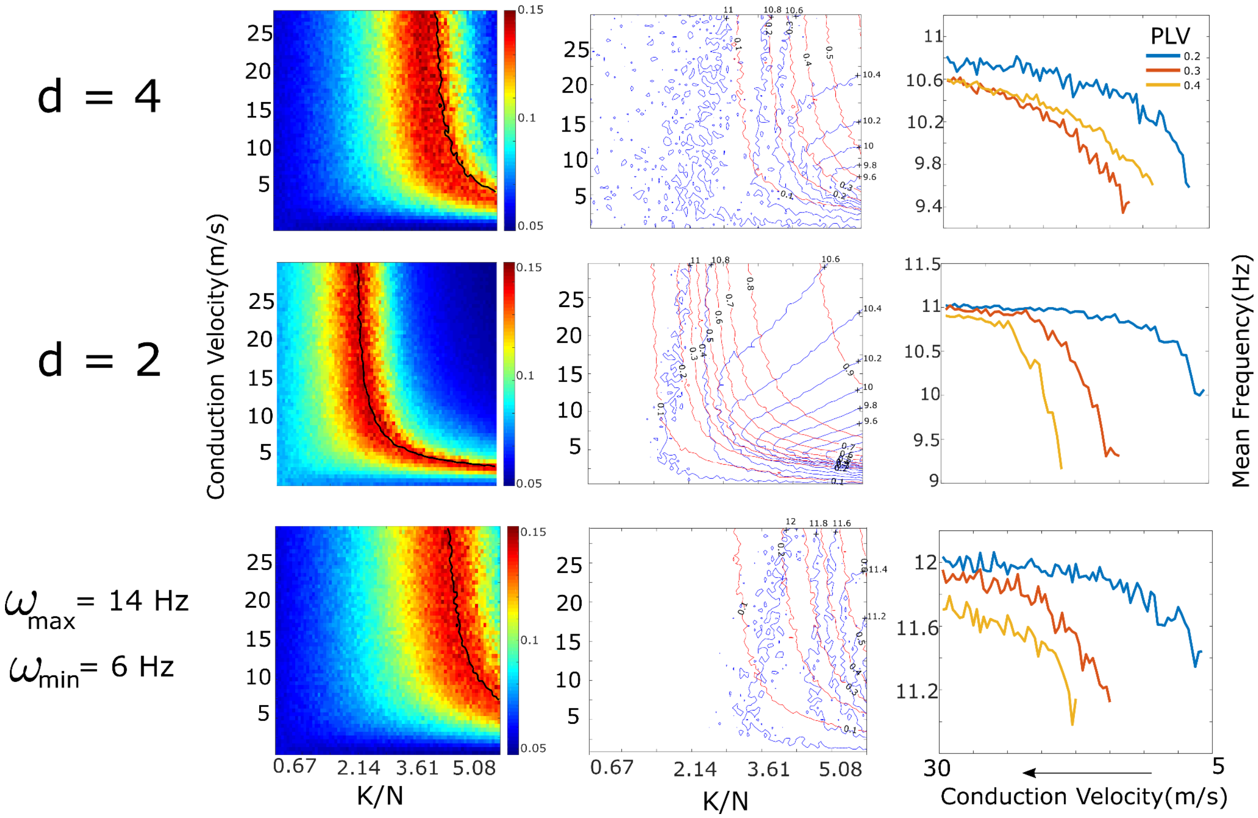


#### 2. Participant information

As outlined in the main text, 200 subjects were randomly sampled from the total 650 subjects for subsequent source localization analysis. Care was taken to ensure similar numbers across age groups and gender. Following are the subject details-

| Younger(18-34) | Age | Sex | Middle-Elderly(35-49) | Age | Sex | Middle-Late(50-64) | Age | Sex | Older(65-88) | Age | Sex |
|---|---|---|---|---|---|---|---|---|---|---|---|
| 110037 | 18 | Male | 221040 | 36 | Male | 420060 | 51 | Male | 510237 | 66 | Male |
| 120061 | 19 | Male | 221324 | 36 | Male | 420202 | 51 | Male | 510226 | 67 | Male |
| 120550 | 19 | Male | 210657 | 37 | Male | 410432 | 52 | Male | 510086 | 65 | Male |
| 110606 | 20 | Male | 222496 | 38 | Male | 420236 | 53 | Male | 610288 | 69 | Male |
| 120212 | 20 | Male | 223085 | 38 | Male | 420396 | 53 | Male | 610496 | 70 | Male |
| 121685 | 20 | Male | 320267 | 38 | Male | 410248 | 54 | Male | 620499 | 71 | Male |
| 110098 | 23 | Male | 320109 | 39 | Male | 410032 | 55 | Male | 610178 | 72 | Male |
| 110101 | 23 | Male | 320297 | 40 | Male | 410040 | 55 | Male | 620567 | 74 | Male |
| 120137 | 18 | Male | 320478 | 40 | Male | 420589 | 52 | Male | 610210 | 75 | Male |
| 122016 | 23 | Male | 222120 | 37 | Male | 410101 | 56 | Male | 610568 | 76 | Male |
| 110033 | 24 | Male | 320022 | 40 | Male | 410226 | 56 | Male | 610052 | 77 | Male |
| 110411 | 25 | Male | 310391 | 41 | Male | 410086 | 57 | Male | 610658 | 78 | Male |
| 120182 | 26 | Male | 320448 | 42 | Male | 420157 | 58 | Male | 621284 | 79 | Male |
| 121144 | 26 | Male | 320461 | 42 | Male | 420198 | 58 | Male | 720103 | 80 | Male |
| 120309 | 27 | Male | 310400 | 43 | Male | 520168 | 59 | Male | 710350 | 81 | Male |
| 120049 | 28 | Male | 310256 | 44 | Male | 510243 | 60 | Male | 720071 | 82 | Male |
| 112141 | 29 | Male | 321107 | 44 | Male | 510258 | 60 | Male | 720407 | 82 | Male |
| 120264 | 28 | Male | 310008 | 45 | Male | 420286 | 56 | Male | 710154 | 83 | Male |
| 120120 | 25 | Male | 321544 | 44 | Male | 410119 | 57 | Male | 720290 | 84 | Male |
| 210174 | 30 | Male | 310331 | 46 | Male | 510551 | 61 | Male | 710131 | 85 | Male |
| 210023 | 31 | Male | 310473 | 46 | Male | 510076 | 62 | Male | 710551 | 85 | Male |
| 220535 | 32 | Male | 310252 | 47 | Male | 520078 | 63 | Male | 721374 | 86 | Male |
| 220223 | 33 | Male | 310142 | 48 | Male | 510329 | 64 | Male | 712027 | 87 | Male |
| 221595 | 33 | Male | 310385 | 48 | Male | 510438 | 62 | Male | 720774 | 87 | Male |
| 210422 | 34 | Male | 420244 | 49 | Male | 420217 | 50 | Male | 510304 | 66 | Female |
| 210617 | 34 | Male | 220234 | 35 | Male | 420226 | 50 | Male | 520134 | 67 | Female |
| 220107 | 34 | Male | 320059 | 48 | Male | 420061 | 57 | Male | 520279 | 68 | Female |
| 221336 | 34 | Male | 220635 | 36 | Female | 510255 | 62 | Male | 610099 | 69 | Female |
| 110182 | 18 | Female | 220198 | 37 | Female | 410323 | 51 | Female | 610625 | 70 | Female |
| 120376 | 18 | Female | 221002 | 37 | Female | 410121 | 52 | Female | 620118 | 71 | Female |
| 120347 | 21 | Female | 310203 | 39 | Female | 410220 | 52 | Female | 610292 | 72 | Female |
| 110056 | 22 | Female | 320575 | 39 | Female | 420204 | 53 | Female | 510355 | 65 | Female |
| 110126 | 22 | Female | 320321 | 40 | Female | 410094 | 54 | Female | 610469 | 73 | Female |
| 120276 | 23 | Female | 310397 | 41 | Female | 410325 | 54 | Female | 620557 | 74 | Female |
| 120727 | 23 | Female | 310410 | 41 | Female | 410222 | 55 | Female | 610392 | 75 | Female |
| 110045 | 24 | Female | 310051 | 42 | Female | 410182 | 53 | Female | 610462 | 76 | Female |
| 110187 | 25 | Female | 310361 | 43 | Female | 410243 | 56 | Female | 610076 | 77 | Female |
| 120065 | 25 | Female | 320661 | 43 | Female | 410084 | 57 | Female | 620354 | 78 | Female |
| 120640 | 26 | Female | 320325 | 44 | Female | 410091 | 57 | Female | 710223 | 79 | Female |
| 120218 | 27 | Female | 310463 | 45 | Female | 410297 | 58 | Female | 720238 | 80 | Female |
| 110069 | 28 | Female | 320576 | 45 | Female | 520197 | 59 | Female | 720622 | 81 | Female |
| 110087 | 28 | Female | 320814 | 43 | Female | 520287 | 59 | Female | 720023 | 82 | Female |
| 210519 | 29 | Female | 310214 | 46 | Female | 510259 | 60 | Female | 710088 | 83 | Female |
| 210148 | 30 | Female | 310086 | 47 | Female | 510433 | 61 | Female | 710342 | 83 | Female |
| 210172 | 31 | Female | 310224 | 47 | Female | 510115 | 62 | Female | 720516 | 84 | Female |
| 220098 | 32 | Female | 312222 | 48 | Female | 510486 | 63 | Female | 710591 | 85 | Female |
| 220511 | 32 | Female | 321899 | 49 | Female | 510323 | 64 | Female | 720400 | 86 | Female |
| 220323 | 33 | Female | 410097 | 49 | Female | 520239 | 65 | Female | 723395 | 86 | Female |
| 210314 | 34 | Female | 420776 | 49 | Female | 420260 | 50 | Female | 721224 | 87 | Female |
| 220203 | 34 | Female | 210124 | 35 | Female | 420435 | 53 | Female | 711035 | 88 | Female |
| 50 | 18 – 34 | Male = 28 | 50 | 36 – 49 | Male = 27 | 50 | 51 – 65 | Male = 28 | 50 | 66 – 88 | Male = 24 |

**3. Generalization of results across different datasets and parcellation schemas**

We replicated our model findings on a separate DTI connectivity matrix as described in Cabral et. al(2014). The connectivity was parcellated according to the Automated Anatomical labelling(AAL) atlas(90 ROIs). Network frequency and phase locking were estimated by varying conduction velocity and global coupling. We observed qualitatively similar results with the AAL atlas. Network frequency reduces as the system of oscillators maintain phase locking at slower conduction velocities.

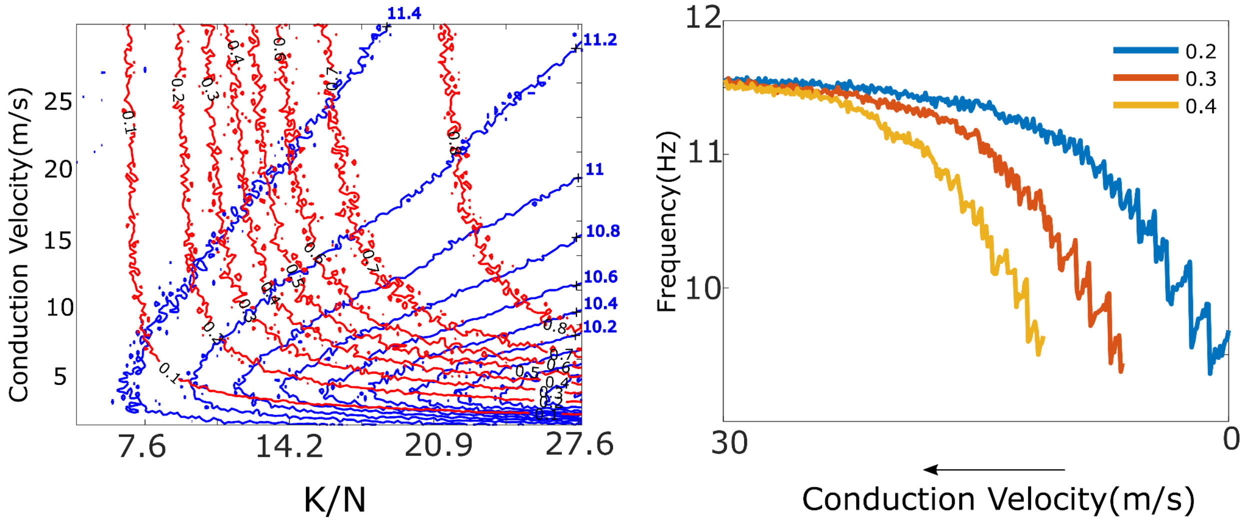


**4. Sensor Space analysis**

We repeated our source-level analysis on MEG sensor level data. Participants possessing a distinct alpha peak in at least half of the magnetometers(N=102) were selected for PLI analysis(587 out of 650 participants).

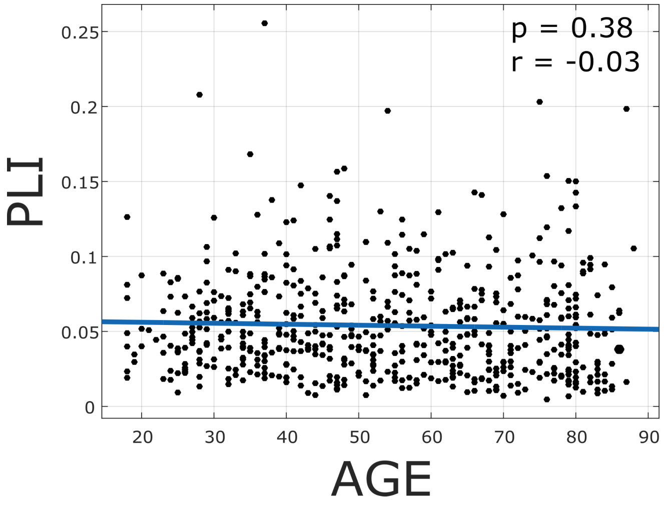

**5. Deriving the macroscopic order parameter for the case of no delays**

The phase dynamics are described by the following equation(**1**)-

$$\dot{\theta}_i = \omega_i + \frac{K}{N}\sum_{j=1}^{N}\sin(\theta_j - \theta_i)$$

The RHS may be written as(**2**)-

$$\frac{1}{N}\sum_{j=1}^{N}\sin(\theta_j - \theta_i) = Im[ze^{-i\theta_i}]$$

We can define a distribution function such that(**3**)-

$$\int_0^{2\pi} f(\omega, \theta, t)d\theta = g(\omega)$$

The distribution function tracks the evolution of the system of oscillators(**4**)-

$$f(\omega, \theta, 0) \rightarrow f(\omega, \theta, t)$$

Since the number of oscillators in the system remains unchanged, the distribution function must obey the following continuity equation(**5**)-

$$\frac{\partial f}{\partial t} + \frac{\partial \dot{\theta} f}{\partial \theta} = 0$$

This implies(6)-

$$\frac{\partial f}{\partial t} + \frac{\partial}{\partial \theta}[(\omega + \frac{K}{2}(ze^{-i\theta} - \tilde{z}e^{i\theta}))f] = 0$$

Since f is a periodic function in θ, it may be written as a Fourier series(**7,8**)-

$$f(\theta, \omega, t) = f(\theta + 2\pi, \omega, t)$$

$$f = \frac{g(\omega)}{2\pi}(1 + \sum_{1}^{+\infty} f_n(\omega, t)e^{in\theta} + \sum_{-\infty}^{-1} \tilde{f}_n(\omega, t)e^{-in\theta})$$

Ott and Antonsen propose the following ansatz(**9**)-

$$f_n(\omega, t) = \alpha^n(\omega, t)$$

Subject to the condition(**10**)-

$$|\alpha(\omega, t)| < 1$$

Inserting equations **9,10** in **7** yields the following(**11**)-

$$\frac{\partial \alpha}{\partial t} + \frac{K}{2}(z\alpha^2 - \widetilde{z}) + i\omega\alpha = 0$$

Inserting 9,10 into 4 yields(**12**)-

$$\widetilde{z} = \int_{-\infty}^{\infty} \alpha(\omega, t) g(\omega) dw$$

**12** may be integrated using the residual theorem by making the substitution(**13**)-

$$z = \overset{*}{\alpha}(\mu - i\gamma, t)$$

Substituting 13 in 11 gives(**14**)-

$$\dot{z} = (i\mu - \gamma)z - \frac{K}{2}(z^2\widetilde{z} - z)$$

For more information about the derivation please refer-

**Edward Ott and Thomas M Antonsen**. Low dimensional behavior of large systems of globally coupled oscillators. *Chaos: An Interdisciplinary Journal of Nonlinear Science*, 18(3):037113, 2008
